# Intrinsic BMP inhibitor Gremlin regulates lung alveolar progenitor cell proliferation and differentiation

**DOI:** 10.1101/2022.07.18.500545

**Authors:** Toyoshi Yanagihara, Quan Zhou, Kazuya Tsubouchi, Spencer Revill, Anmar Ayoub, Sy Giin Chong, Anna Dvorkin-Gheva, Kjetil Ask, Wei Shi, Martin RJ Kolb

**Affiliations:** Firestone Institute for Respiratory Health, Research Institute at St Joseph’s Healthcare, Department of Medicine, McMaster University, Hamilton, ON, Canada; Department of Respiratory Medicine, Graduate School of Medical Sciences, Kyushu University, Fukuoka, Japan; McMaster Immunology Research Centre, Department of Medicine, McMaster University, Hamilton, ON, Canada; Department of Surgery, Children’s Hospital Los Angeles, Keck School of Medicine, University of Southern California, Los Angeles, California

**Keywords:** Gremlin, Bone morphogenic protein, alveolar epithelial cell, alveolar progenitor cell

## Abstract

Type 1 alveolar epithelial cells (AT1s) and type 2 alveolar epithelial cells (AT2s) regulate the structural integrity and function of alveoli. AT1s mediate gas exchange, whereas AT2s serve multiple functions, including surfactant secretion and alveolar repair through proliferation and differentiation into AT1s as progenitors. However, mechanisms regulating AT2 proliferation and differentiation remain unclear. Here we demonstrate that Gremlin, an intrinsic inhibitor of bone morphogenetic protein (BMP), induces AT2 proliferation and differentiation. Transient overexpression of Gremlin in rat lungs by adenovirus vector delivery suppressed BMP signaling, induced proliferation of AT2s and the production of Bmp2, which in turn led to the recovery of BMP signaling and induced AT2 differentiation into AT1s. Gremlin was upregulated in a bleomycin-induced lung injury model. TGF-*β* and IL-1*β* induced Gremlin expression in fibroblasts. Taken together, our findings implicate that Gremlin expression during lung injury leads to precisely timed inhibition of BMP signaling and activates AT2s, leading to alveolar repair.

## Introduction

The lung is a complex organ encompassing air-conducting tubes terminating in millions of air-exchanging units called alveoli. The alveolar epithelium is composed of two distinct cell types: type 1 alveolar epithelial cells (AT1s) and type 2 alveolar epithelial cells (AT2s). AT1s are thin squamous cells that cover about 95% of the internal surface of the lung and are essential for gas exchange between the air in the alveoli and blood in the capillaries (1). AT2s, are cuboidal epithelial cells that maintain the surfactant biosynthetic function (2) but also serve as alveolar progenitor cells capable of self-renewal and differentiation into AT1s to repair the damaged epithelium (3)(4). However, the molecular mechanisms underlying AT2 proliferation and AT1 differentiation are not fully understood, but recent evidence suggests the involvement of Bone Morphogenetic Protein (BMP) antagonists (15).

Gremlin is an extracellular glycoprotein with strong antagonistic activity on BMPs that are highly expressed in the lungs (5)(8). Studies in mice with Gremlin null mutation have shown that Gremlin plays an essential role in lung development, particularly lung branching (6)(7). However, its physiological role in the lung after development is still unknown. It has been reported that Gremlin may be involved in the pathogenesis of some lung diseases, such as pulmonary hypertension (8) and idiopathic pulmonary fibrosis (IPF), yet the mechanism behind this is unclear (9)(10)(11). Interestingly, we find that transient Gremlin overexpression along with the dynamic change in lung BMP signaling induces AT2 proliferation, followed by subsequent AT1 differentiation and fibrosis resolution. In bleomycin-induced lung injury, Gremlin is overexpressed. Transforming growth factor-beta (TGF-*β*) or pro-inflammatory cytokines, such as interleukin-1*β* (IL-1*β*), also induce Gremlin expression *in vitro*. This suggests that Gremlin plays a role in lung injury and repair through altering BMP signaling and promoting AT2 proliferation and subsequent AT1 differentiation.

## Materials and Methods

### Animal Experiments

Transient Gremlin overexpression was induced by an adenoviral vector encoding Gremlin 1 (hereafter designated as AdGrem), and transient TGF-*β*1 overexpression was induced by an adenoviral vector encoding an active form of TGF-*β*1 (hereafter designated as AdTGF-*β*1). Female Sprague-Dawley rats (250–300 g; Charles River, Wilmington, MA) received 1.0 × 10^8^ PFU of AdGrem or 5.0 × 10^8^ PFU of AdTGF-*β*1 by single intratracheal instillation under isoflurane anesthesia on day 0 and sacrificed on days 7, 14, and 28. Control rats received an empty vector construct (AdDL). For the bleomycin-induced lung injury model, rats received 2.5 U/kg of bleomycin by single intratracheal instillation. Control rats received phosphate-buffered saline (PBS). Lungs were harvested and either fixed in 10% formalin for histology (left lobes) or flash-frozen in liquid nitrogen for protein and RNA analysis (right lobes). All animal work was conducted under the guidelines from the Canadian Council on Animal Care and approved by the Animal Research Ethics Board of McMaster University under protocol #17-07-31.

### Human samples

All tissues were collected with patient consent in compliance with the Research Ethics Board of St. Joseph’s Healthcare Hamilton. Hamilton Integrated Research Ethics Board (HIREB #00-1839) approval was obtained prior to beginning the study.

### Antibodies and reagents

Antibodies are GAPDH (#5174, Cell signaling technology)., Gremlin (#4383, Cell Signaling Technology), Phospho-Smad1/5 (#9516S, Cell Signaling Technology), alpha-Tubulin (#2144S, Cell Signaling Technology), PCNA (#2586S, Cell Signaling Technology), Aquaporin 5 (ab92320, abcam), Axin2 (ab109307, abcam), Vimentin (ab16700, abcam), BMP-2/4 (sc-137087, Santa Cruz for western blot), BMP2 (PA578874, Thermofisher for immunofluorescence staining), Anti-rabbit HRP linked IgG (#7074, Cell Signaling Technology), Anti-mouse IgG HRP-linked Antibody (#7076, Cell Signaling Technology). For fluorescence microscopy, we used goat or donkey secondary antibodies conjugated with Alexa Fluor-488 and Alexa Fluor-594 (Abcam) for fluorescence microscopy. Human TGF-β1 (240-B) and IL-1*β* (201-LB-005) were purchased from R&D systems.

### Tissue microarray and RNA-Chromogenic in situ hybridization (CISH)

Formalin-fixed paraffin-embedded (FFPE) human lung tissues were obtained from the biobank for lung diseases at St. Joseph’s Hospital in Hamilton, Ontario. IPF (*n* = 24) cases and controls (non-involved tissues from lung cancer cases) (*n* = 16) were used. Upon confirmation of a positive diagnosis, patient H&E slides were made and scanned using the Olympus VS120 Slide Scanner. Areas of fibrotic or non-fibrotic were selected with the help of a trained molecular pathologist, and 1 mm diameter cores were selected to be placed into a tissue microarray (TMA) block, 4 cores per case were picked for each specified area, using TMA Master II by 3DHISTECH.Ltd machine.

The cellular data was acquired through the HALO® Image Analysis Platform by Indica Labs. The quantitative tissue analysis was performed using HALO®’s ISH (In Situ Hybridization) module in combination with HALO®’s TMA (Tissue Microarray) module. This technology utilizes digitized histology slides to detect and quantify the total number of cells present in each tissue core of the TMA, the number of nucleic acid probes, as well as the percentage of cells that contain those nucleic acid probes. For this experiment, HALO® was used to quantify the number of *GREM1*-positive cells and probe negative cells found in each core. An RNA probe for *GREM1* (312838) was purchased from Advanced Cell Diagnostics.

### Cell preparation and culture

Primary human fibroblasts were obtained from ATCC (PCS-201-013) and grown in Dulbecco’s modified Eagle Medium (DMEM) (Biowhittaker® Reagents, Lonza) supplemented with 10% fetal bovine serum (Gibco) and 1% penicillin-streptomycin (Gibco, Life Technologies) and used at passages between P2 and P5. Cells were cultured at 37°C, 5% CO_2_.

### Histology

Rat lungs of left lobes were fixed by intratracheal instillation of 10% neutral-buffered formalin at a pressure of 20 cm H_2_O. Paraffin sections were cut at 4 μm and processed in-house at the core histology facility at McMaster University (Hamilton, ON, Canada). Tissue slides were stained with Masson Trichrome. Picture acquisition of Masson Trichrome staining was performed using an Automatic slide scanner microscope (Olympus VS 120-L).

### Immunofluorescence

Immunostaining was performed on formalin-fixed rat lung tissue sections. Briefly, following deparaffinization, antigen retrieval with citric acid, and saturation of nonspecific sites with 10% horse serum/PBS for 30 min, lung sections were incubated with primary antibodies overnight in a humidified chamber at 4°C. Conjugated secondary antibodies were used at a dilution of 1:500. Slides were mounted in Prolong-gold with DAPI (ProLong® Gold antifade reagent with DAPI, Life technologies, P36931).

### Western blotting

Crushed lungs were homogenized using a mechanical homogenizer (Omni International, Waterbury CT) or cultured cells were lysed in 1× lysis buffer (9803S, NEB) (20 mM Tris-HCl buffer (pH 7.5) containing 1% Triton X-100, 150 mM NaC1, 1 mM EDTA, 1 mM EGTA, 2.5 mM sodium pyrophosphate, 1 mM *β*-glycerophosphate, 1 mM Na3VO4) supplemented with complete protease inhibitors (Roche) and the collected supernatant was used for western blotting. Total protein from lung homogenate (30–60 µg) or cells (20 µg) was separated on a 6–12% SDS Polyacrylamide Electrophoresis gels based on the molecular weight. Proteins were transferred to a PVDF membrane (Bio-Rad Laboratories,1620177, Hercules, CA) using a wet transfer apparatus and blocked at room temperature for 30 min using 5% skim milk. Protein detection was performed using Clarity(tm) Western ECL Substrate (Bio-Rad, 1705060) and analyzed in a ChemiDoc XRS Imaging System (Bio-Rad Laboratories). The signals were measured using ImageJ public-domain software.

### Isolation of mRNA and gene expression

Total RNA was extracted from either crushed frozen lung tissue or cultured cells with TRIzol® reagent (Thermo fisher scientific, 15596026) according to the manufacturer’s recommendations. RNA was reverse transcribed using qScript cDNA SuperMix (Quanta Bioscience, 95048-025, Gaithersburg, MD). The cDNA was amplified using a Fast 7500 real-time PCR system (AB Applied Biosystems) using TaqMan® Universal PCR Master Mix and predesigned primer pairs (Thermo Fisher Scientific) for rat *Gapdh* (Rn01775763_g1), rat *Grem1*(Rn01509832_m1), rat *Sftpc*(Rn00569225_m1), rat *Aqp5*(Rn00562837_m1), human *GREM1*(Hs01879841_s1), and human *GAPDH*(4333764E).

### Single-cell RNA sequencing

Data preprocessed using the Cell Ranger pipeline (10x Genomics) were obtained from GSE122960. All 8 available samples from donors and 4 available samples from IPF patients were selected from the dataset, downloaded and used for further analysis. Post-processing was performed using the same pipeline as in the source paper (12) with *seurat* (13) package in R. Visualizations of both violin plots were created using *seurat*. Cell populations were defined using the genes reported to be differentially expressed between the cell populations defined in the source publication (12).

### Statistical Analysis

The Student’s two-tailed unpaired t-test was used to compare the two groups. Statistical analysis between multiple groups with one control group was performed by one-way analysis of variance (ANOVA) followed by Dunnett’s multiple comparison test with the use of GraphPad Prism 8 (GraphPad Software Inc.). A *p*-value less than 0.05 was considered significant.

## Results

### Upregulated Gremlin expression in lung sections from IPF patients

First, we confirmed Gremlin upregulation in IPF at both mRNA and protein levels. We investigated Gremlin expression in lung sections. Gremlin was upregulated in the fibrotic area of IPF lungs (Figures 1A and 1B). This Gremlin upregulation in IPF is consistent with the results in a previous study (9). Immunofluorescence staining showed that Gremlin expression was co-localized with fibroblasts (vimentin-positive cells), myofibroblasts (*α*SMA-positive cells), and macrophages (CD68-positive cells) (Figures 1C and 1D), suggesting that fibroblasts, myofibroblasts, and macrophages and are the sources of Gremlin in fibrotic lungs.

**Figure 1.**
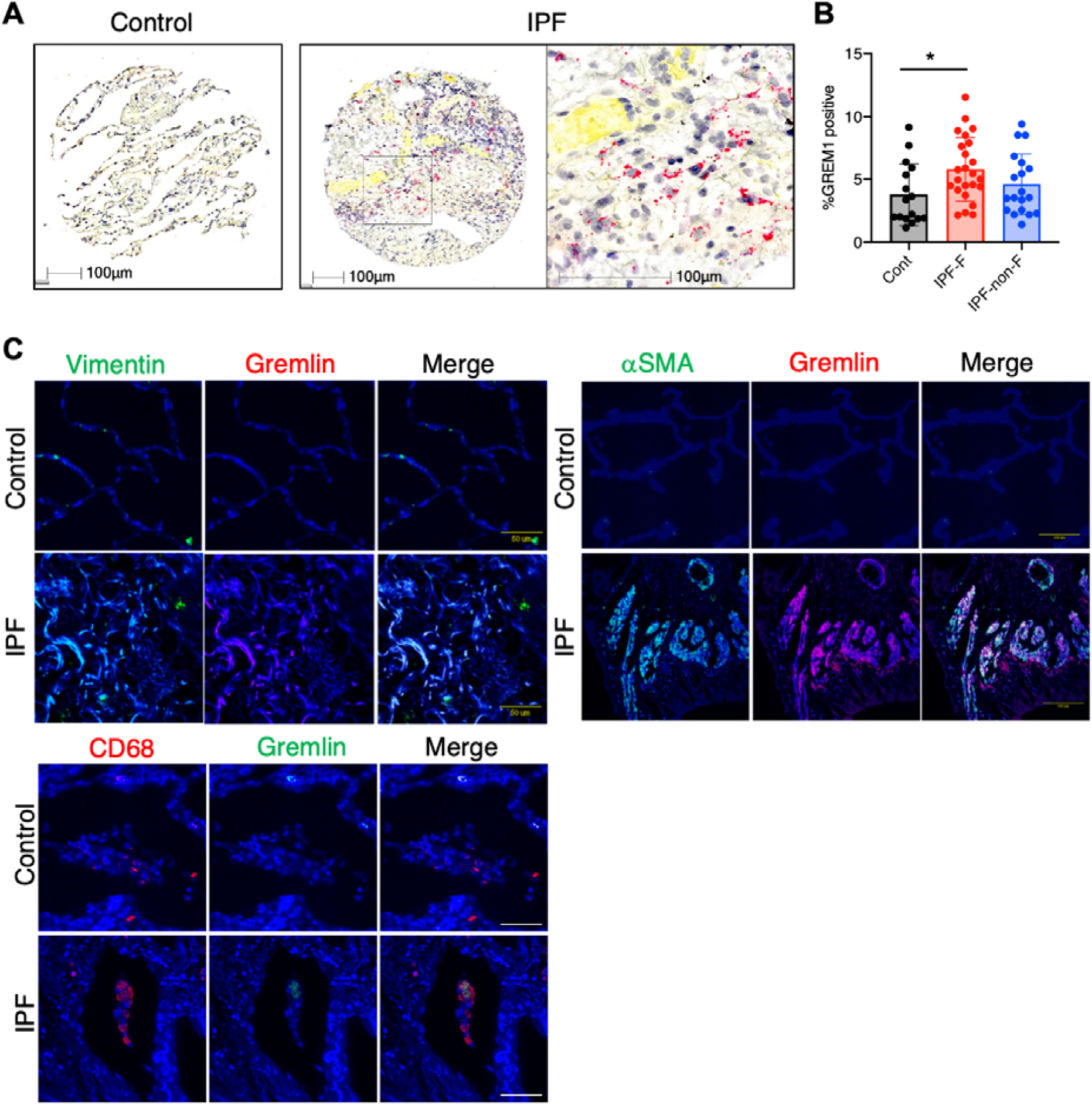
Upregulated Gremlin expression in idiopathic pulmonary fibrosis. (A) Representative images and quantification (B) of RNA-chromogenic *in situ* hybridization for *GREM1* (red) in human lung sections from control (*n* = 16) and idiopathic pulmonary fibrosis (IPF) (fibrotic area (IPF-F): *n* = 24, non-fibrotic area (IPF-non-F): *n* = 19). (C)(D) Immunofluorescence staining of Gremlin, Vimentin, *α*SMA, CD68, and DAPI on lung sections from healthy control and a patient with IPF. \**p* < 0.05.

### Marked overgrowth of AT2s in rat lung overexpressing Gremlin

Gremlin overexpression in rat lung was achieved by intra-tracheal delivery of adenoviral vectors. Masson and Trichrome staining analyses showed alveolar wall thickening with prominent epithelial cell expansion in AdGrem-treated lungs on day 7 (Figure 2A), while no cellular changes in AdDL-treated lungs were observed, consistent with our previous study (14). Immunofluorescent staining analysis revealed that the ratio of AT2 (surfactant protein-C (SP-C) positive cells) to AT1 (Aquaporin 5 (Aqp5) positive cells) was markedly increased in AdGrem-treated lung (Figure 2B). Western blot and qPCR analysis confirmed the change in SP-C/Aqp5 ratio in AdGrem-treated lungs. (Figures 2C, 2D, and 2E). The percentage of PCNA^+^ AT2 was greatly increased in AdGrem-treated lungs (AdDL vs. AdGrem: 13.1 ± 3.5% vs. 78.6 ± 6.7%), suggesting the proportional change of AT2/AT1 was due to AT2 proliferation. Recently, a study identified that the Wnt pathway gene Axin2+ AT2s have alveolar progenitor cell activity (15). Immunofluorescent staining revealed Axin2+ AT2s were increased in AdGrem-treated lungs (Figure 2H); western blot analysis confirmed increased levels of Axin2 in AdGrem-treated lungs (Figure 2I). Together, this suggests that Gremlin stimulates AT2 with an alveolar progenitor cell function. Thus, overexpression of Gremlin in the lung resulted in the marked proliferation of AT2, which might have an alveolar progenitor cell activity.

**Figure 2.**
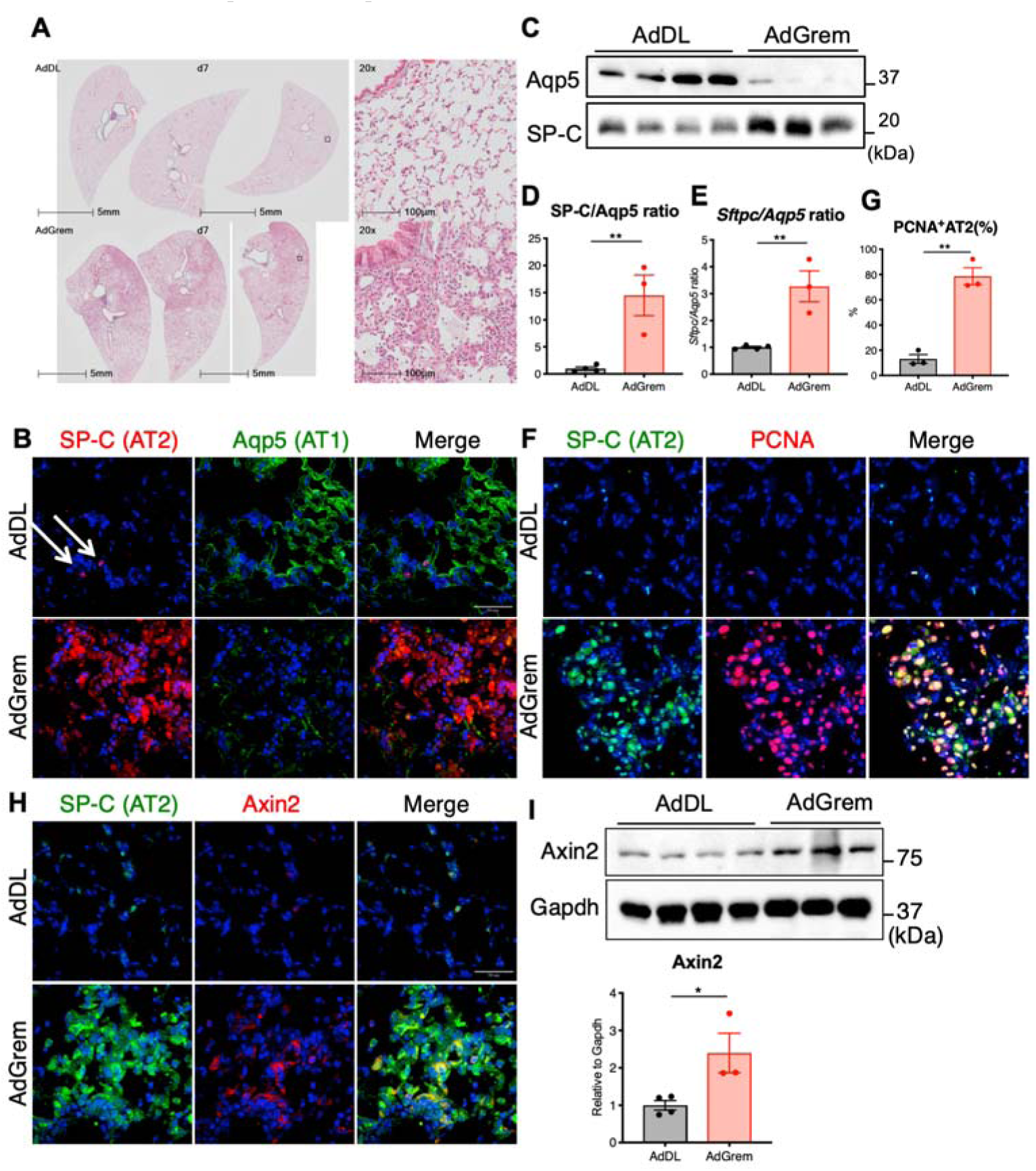
Marked overgrowth of type 2 alveolar epithelial cells (AT2) in AdGremlin-treated rat lung on day 7. (A) Representative lung sections stained by Masson trichrome imaged by Slide Scanner from AdDL and AdGremlin (AdGrem) on day 7. (B) Immunofluorescence staining (IF) of surfactant protein-C (SP-C) and Aquaporin 5 (Aqp5), or SP-C and Axin2 (F), and DAPI on the lung sections from rats treated with AdDL and AdGrem on day 7. Arrows indicate positive signals for SP-C. (C) Western blot analysis and quantification (D) of SP-C and Aqp5 in whole lung lysates of AdDL and AdGrem on day 7. (E) qPCR analysis of *Sftpc*/*Aqp5* ratio in whole lung tissues. (F) IF of SP-C, PCNA, and DAPI on the lung sections from rats treated with AdDL and AdGrem. (G) Quantification of PCNA positive proportion in AT2 (SP-C^+^ cells) on the lung sections from rats treated with AdDL and AdGrem. (I) Western blot analysis and quantification of Axin2 in whole lung lysates of AdDL and AdGrem. Gapdh was used as a loading control. AdDL; *n* = 4, AdGrem; *n* = 3. Data are expressed as means ± SEM. \*\**p* < 0.01, \**p* < 0.05.

### Decreased BMP signaling with elevated levels of Bmp2 in AdGrem-treated rat lung

To evaluate the effect of Gremlin on BMP signaling in the lung, phosphorylation of Smad1/5 (pSMAD1/5), a marker of intracellular BMP signal activation, was measured. On day 7 when Gremlin expression was upregulated, pSMAD1/5 level was down-regulated in AdGrem-treated lungs (Figures 3A and 3B), verifying the proof of concept of BMP signaling suppression. Surprisingly, however, the levels of Bmp2, a BMP ligand itself, were greatly increased in AdGrem-treated lungs, suggesting a compensatory mechanism (Figures 3A and 3B).

**Figure 3.**
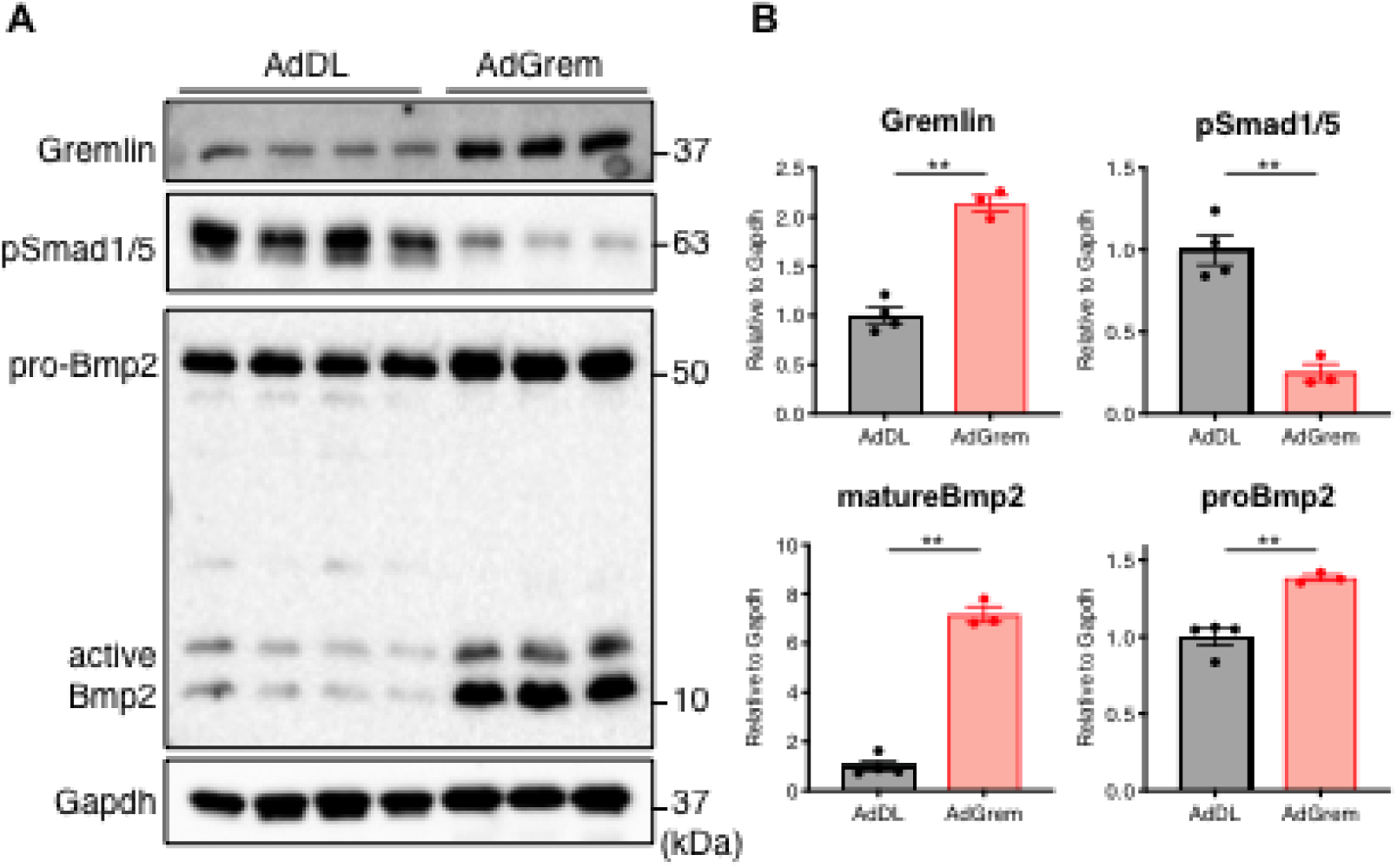
Decreased BMP signaling with elevated levels of Bmp2 in AdGrem-treated rat lung on day 7. (A) Western blot analysis and quantification (B) of Gremlin, phospho-Smad1/5 (pSmad1/5), and Bmp2 in whole lung lysates of AdDL and AdGremlin (AdGrem) on day 7. Gapdh was used as a loading control. AdDL; *n* = 4, AdGrem; *n* = 3. Data are expressed as means ± SEM. **p < 0.01, *p < 0.05.

### AT2s are the main sources of Bmp2 in AdGrem-treated lung

To understand the unexpected result of increased levels of Bmp2, we analyzed previously published and publicly available single-cell RNA sequencing (scRNA-seq) data from human lung tissues (12) to see the sources of BMPs in the lung. scRNA-seq showed that some AT2s produce Bmp2 and Bmp4 (Figure 4A). Immunostaining revealed Bmp2 expression was mainly co-localized with SP-C (AT2s) expression in AdDL-treated lungs; this expression and co-localization was greatly increased in AdGrem-treated lungs (Figure 4B). This suggests the increased expression of the Bmp2 ligand shown in the AdGrem-treated lungs is sourced from proliferating AT2s.

**Figure 4.**
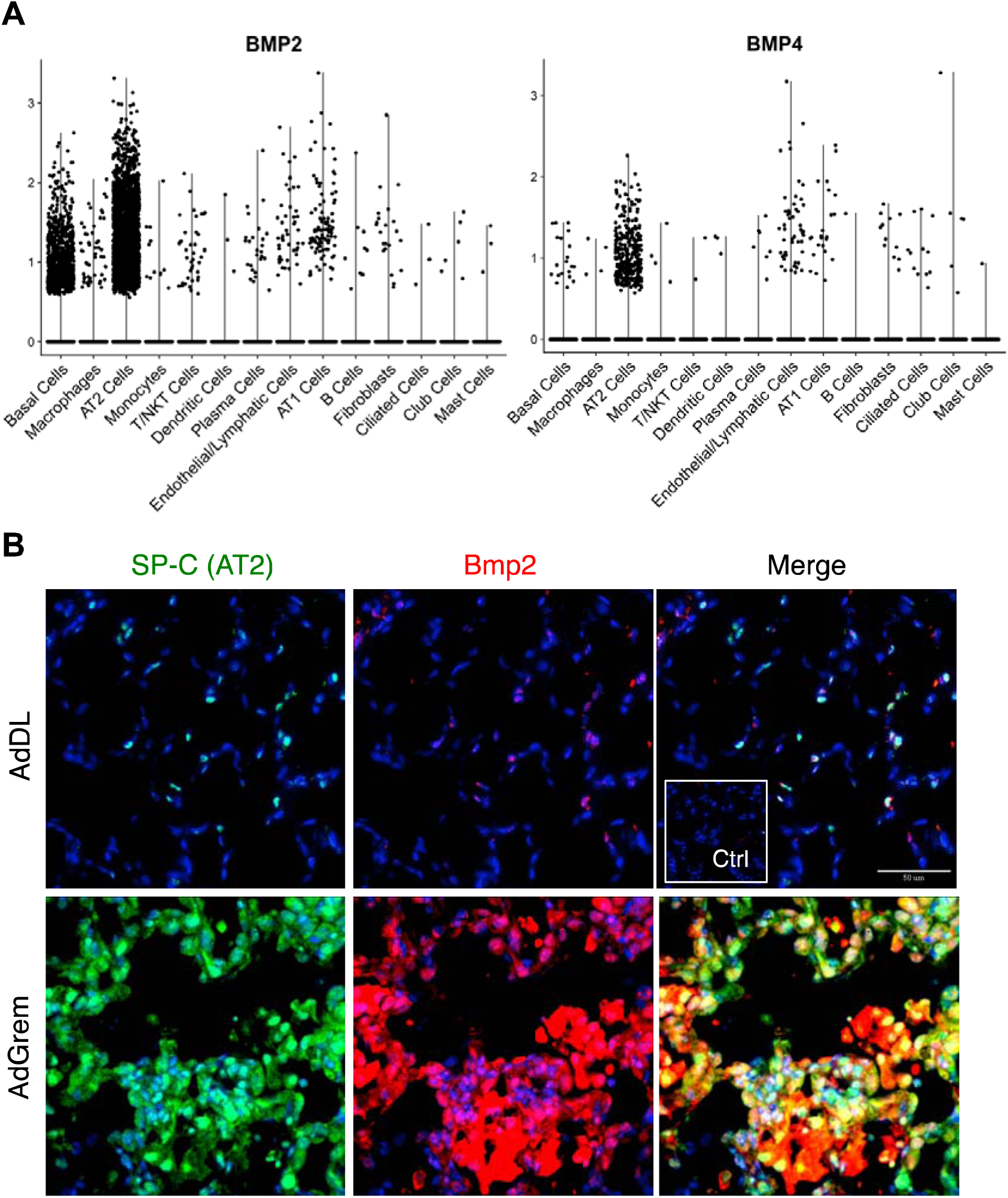
Type 2 alveolar epithelial cells (AT2s) were the main sources of Bmp2/Bmp4. (A) Expression levels of *BMP2* or *BMP4* across cell populations in human lung tissues. single-cell RNA sequence was performed on samples obtained from 8 lung biopsies from transplant donors and 4 lung explants from transplant recipients with IPF. (B) Immunofluorescence staining of surfactant protein-C (SP-C), Bmp2, and DAPI on the lung sections from rats treated with AdDL and AdGremlin (AdGrem) on day 7.

### Dynamics of AT2/AT1 ratio and BMP signaling in AdGrem-treated lung

Next, the relationship between epithelial phenotype (AT2/AT1 ratio) and BMP signaling in AdGrem-treated lungs was explored. Recently, Chung *et al*. reported that BMP signaling declined transiently in AT2 cells in a post-pneumonectomy lung in vivo, associated with up-regulation of BMP antagonists, and is restored during differentiation of AT2 to AT1 (16). We previously showed that the transgenic product Gremlin disappears by day 14 in this adenovirus gene transfer (14).

Levels of Gremlin and Bmp2 in AdGrem-treated lungs were increased on day 7 and normalized by day 28, while levels of BMP signaling (pSmad1/5) were decreased on day 7 and approached normalization on day 28 (Figures 5A and 5B). In tandem with these changes, the SP-C/Aqp5 ratio was also normalized in a similar timeframe; a large shift towards SP-C positivity on day 7 with increased progenitor/proliferation markers which normalized on day 28. (Figures 5A and 5B). Immunostaining revealed decreased SP-C positive cells with the increase of Aqp5 positive cells in AdGrem-treated lungs on day 14 compared to that on day 7 (Figure 5C). This, in combination with Chung *et al*. (16), suggests a relationship between the level of BMP signaling present and the level of AT2 progenitor activity. Furthermore, this suggests that Gremlin overexpression suppresses BMP signaling in AT2, resulting in the proliferation of AT2 and production of Bmp2, which in turn reverses BMP signaling in AT2 and induces AT1 differentiation (Figure 5D).

**Figure 5.**
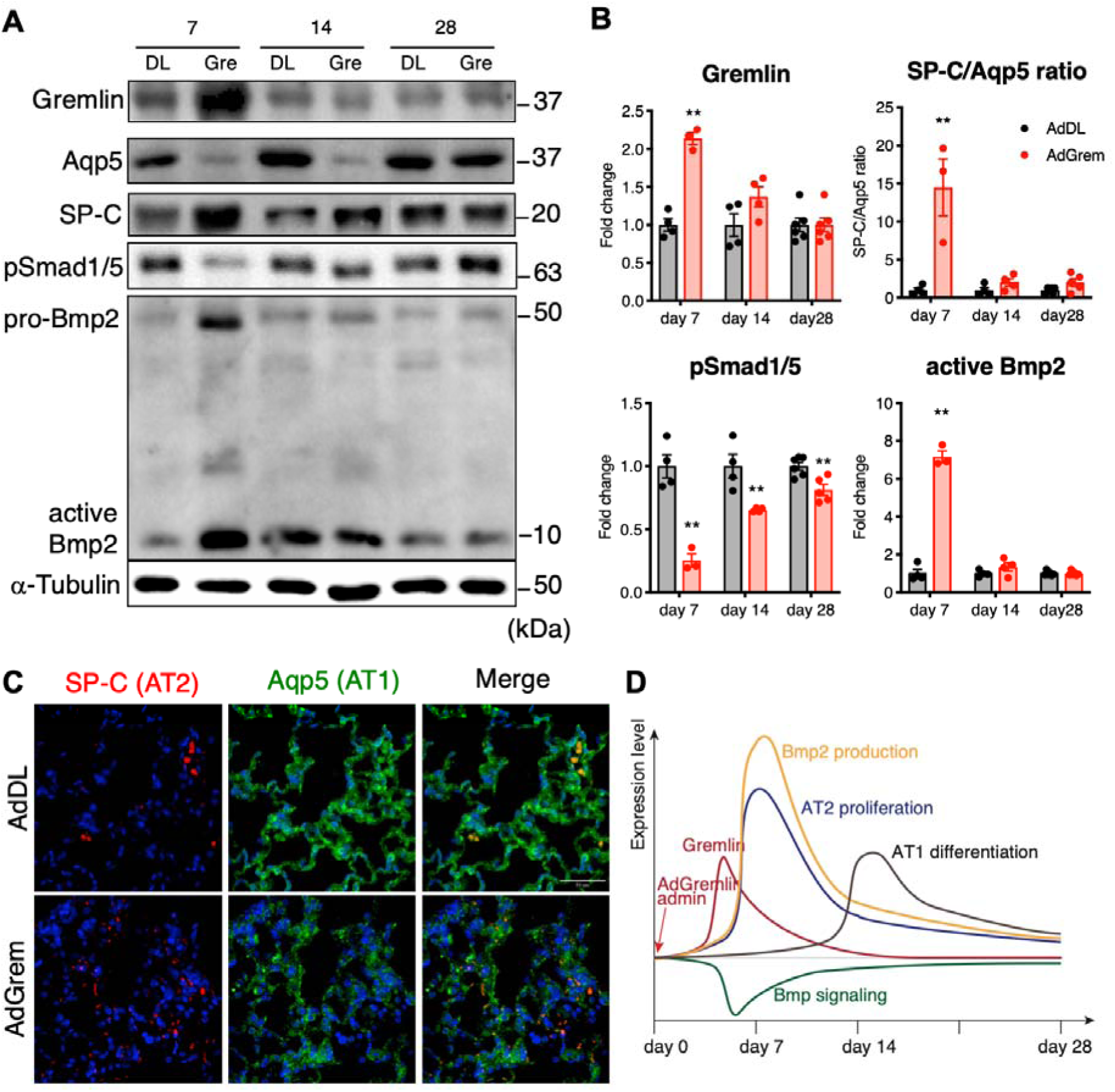
Dynamics of type 2/type 1 alveolar epithelial cell (AT2/AT1) ratio in relation to BMP signaling in AdGrem-treated lung. (A) Western blot analysis of Gremlin, Aquaporin 5 (Aqp5), surfactant protein-C (SP-C), phospho-Smad1/5 (pSmad1/5), Bmp2, and *α*Tubulin in whole lung lysates of AdDL and AdGremlin (AdGrem) on days 7, 14, 28. *α*Tubulin was used as a loading control. (B) Quantification of Western blot intensity in whole lung lysates of AdDL and AdGrem. day 7: AdDL; *n* = 4, AdGrem; *n* = 3. day 14: AdDL; *n* = 4, AdGrem; *n* = 4. day 28: AdDL; *n* = 6, AdGrem; *n* = 5. Data are expressed as means ± SEM. \*\**p* < 0.01, \**p* < 0.05. (C) Immunofluorescence staining of surfactant protein-C (SP-C), Aquaporin 5 (Aqp5), and DAPI on the lung sections from rats treated with AdDL and AdGrem on day 14. (D) Diagram showing the dynamics of Gremlin, BMP signaling, and type 2/type 1 alveolar epithelial cell (AT2/AT1) ratio in AdGrem-treated rat lungs.

### Gremlin expression was upregulated in a bleomycin-induced lung injury model

To investigate the gremlin expression dynamics during lung injury and repair, we next administered bleomycin to rats intratracheally. Intratracheal bleomycin installation is a commonly used rodent model of lung injury and repair (17). Bleomycin initially damages the alveolar epithelial cells, followed by inflammation and fibrosis, with eventual resolution after a single dose of damage. (17). TGF-*β* was upregulated on days 7, 14, and 28, with a peak on day 7 (Figures 6A and 6B). The level of Gremlin was also upregulated (2.6 times compared to the vehicle group) on day 7 but was returned to normal after day 14 (Figures 6A and 6B). Previously published scRNA-seq showed that Gremlin is expressed in several cell types in the lung, including alveolar epithelium, fibroblasts, and macrophages (18). Indeed, fibroblasts (vimentin-positive cells) and macrophages (CD68-positive cells) highly express Gremlin in the fibrotic phase of bleomycin-treated lungs (Figures 6C and 6D), which are consistent with the results of immunofluorescent staining in human lungs (Figures 1C and 1D).

**Figure 6.**
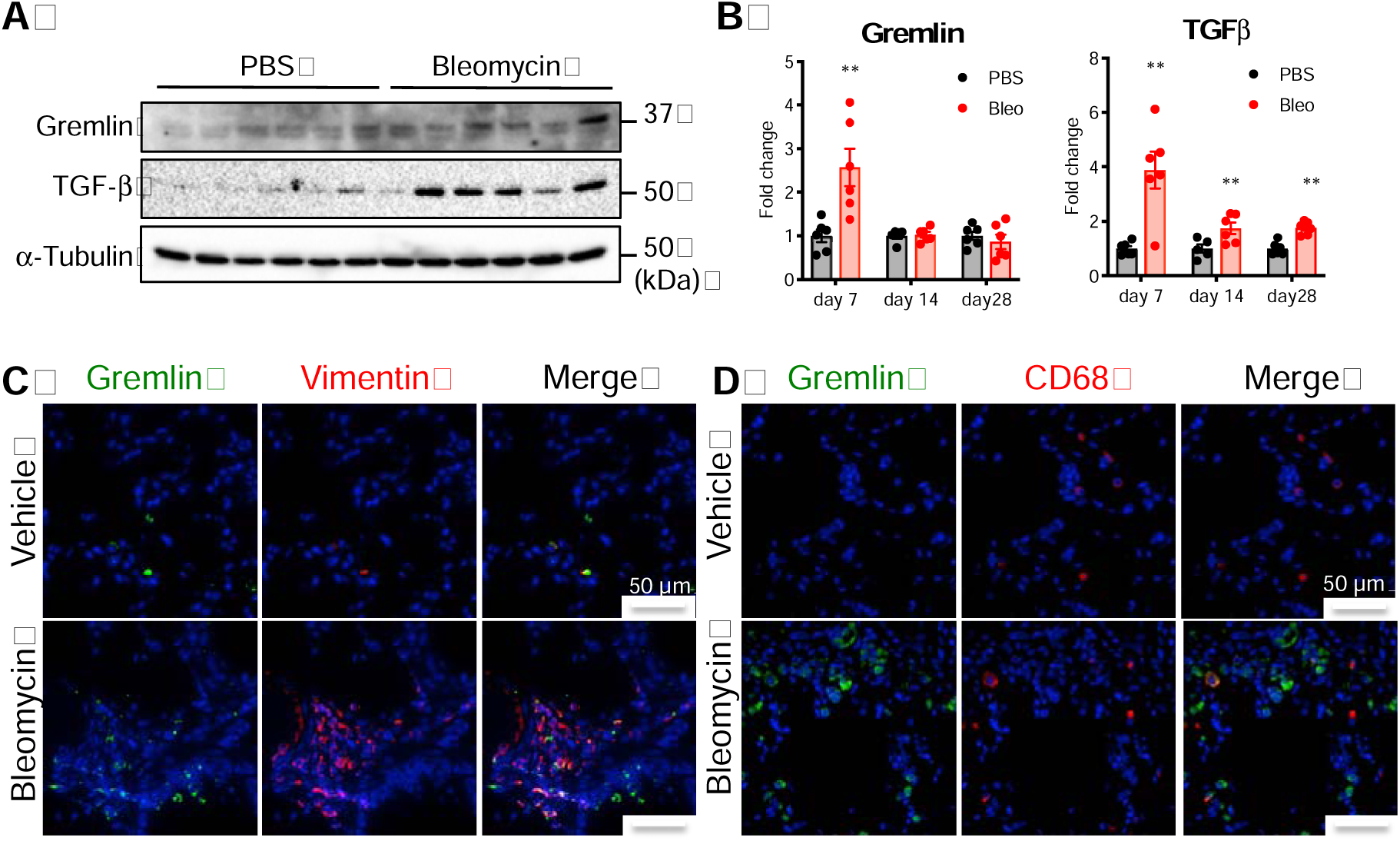
Gremlin expression in bleomycin-induced lung injury model. (A) Western blot analysis of Gremlin, TGF-*β*, and *α*Tubulin in whole lung lysates from rats treated with bleomycin or PBS (Vehicle) on day 7. *α*Tubulin was used as a loading control. (B) Quantification of Western blot intensity in whole lung lysates from rats treated with bleomycin or PBS on days 7, 14, 28. Day 7: PBS; *n* = 6, Bleomycin; *n* = 6. day 14: PBS; *n* = 5, Bleomycin; *n* = 6. day 28: PBS; *n* = 6, Bleomycin; *n* = 6. Data are expressed as means ± SEM. \*\**p* < 0.01. Immunofluorescence staining of Gremlin with Vimentin (C) or CD68 (D), and DAPI on the lung sections from rats treated with Bleomycin or PBS (Vehicle) on day 7.

### Gremlin regulation by TGF*β*, IL-1*β*, and stiffness around cells

To understand the regulatory mechanism for Gremlin, we evaluated factors that can affect Gremlin expression. As TGF-*β* was reported to induce Gremlin expression in human proximal tubule kidney epithelial cells (HK-2) (19), we treated human fibroblasts with recombinant TGF-*β*1 *in vitro*. Recombinant TGF-*β*1 induced Gremlin expression in human fibroblasts in a dose-dependent manner (Figures 7A and 7B), like previously published studies (19). The Gremlin proximal promoter contains two CpG islands with motifs related to NF*κ*B (19), suggesting pro-inflammatory cytokines can also induce Gremlin expression. Indeed, 10 pg/ml of recombinant IL-1*β* induced Gremlin expression in human fibroblasts *in vitro* (Figure 7C). These results suggest that upregulated TGF*β* and pro-inflammatory cytokine IL-1*β* induces Gremlin expression at the damaged alveolar site during lung injury.

**Figure 7.**
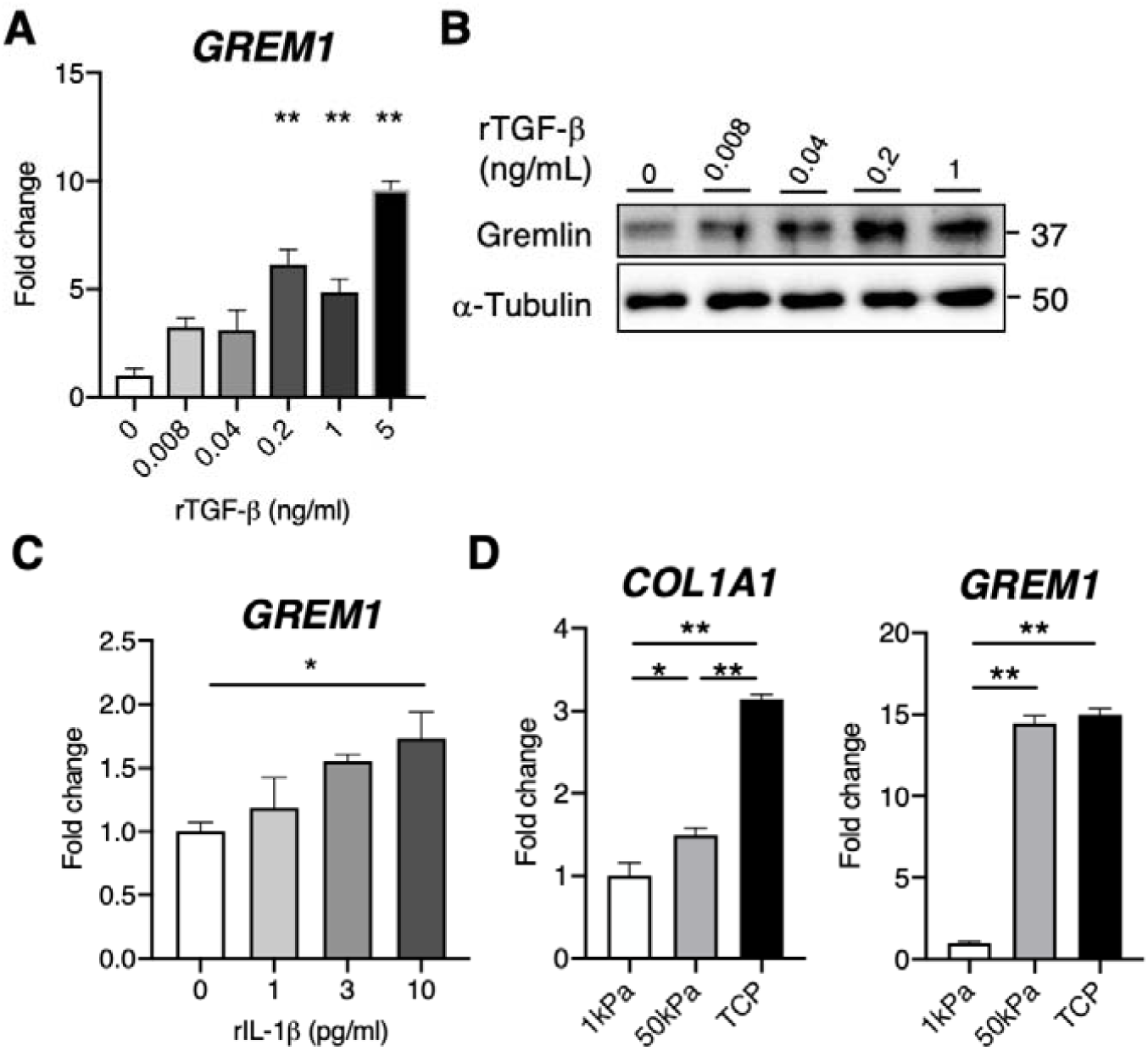
Gremlin regulation by TGF*β*, IL-1*β*, and stiffness. mRNA expression of *GREM1*in human lung fibroblasts treated with recombinant TGF-*β* (A) and (C) IL-1*β* for 24 hrs under serum-free conditions (*n* = 3). *GAPDH* was used as a loading control. (B) Western blot analysis of Gremlin in human lung fibroblasts treated with recombinant TGF-*β* for 24 hrs under serum-free condition. (D) mRNA expression of *COL1A1* and *GREM1* in human lung fibroblasts cultured on plates of differing stiffness. TCP; tissue culture plate. *GAPDH* was used as a loading control. Data are expressed as means ± SEM. \*\**p* < 0.01, \**p* < 0.05.

Gremlin was upregulated in fibroblasts in the fibrotic lung (9). To determine whether lung stiffness and attribute of the fibrotic lung environment induces Gremlin expression in fibroblasts, we cultured fibroblasts on matrices of differing stiffnesses (1 kPa, 50 kPa, and tissue culture plate [TCP]). The physiological stiffness of the lungs is between 0.2 and 2 kPa, whereas the pathological stiffness of the IPF lung is a mean of 16 kPa, up to around 200 kPa (20)(21)(22). TCP is a very rigid matrix, similar to tendon or bone, with a tensile strength of 10^6^ kPa (23). The level of *COL1A1*, one of the fibrotic markers, was highest in TCP, followed by 50 kPa compared to 1 kPa (Figure 7D), suggesting stiffness around the fibroblasts induces a fibrotic phenotype, similar to other pro-fibrotic markers, such as *CTGF* (24). The level of *GREM1* was greatly upregulated on 50 kPa compared to 1 kPa, but no difference between 50 kPa and TCP (Figure 7E). This result indicates that stiffness up to 50 kPa increased Gremlin expression in fibroblasts, while the effect reaches its upper limit at 50 kPa.

## Discussion

We previously reported that transient Gremlin overexpression in the lungs through an adenovirus encoding vector resulted in epithelial cell activation and reversible fibrosis (14). Reversible fibrosis and wound healing have been postulated to be the two sides of the same coin In conjunction with a paper by Chung *et al*. (16), we suggest that BMP signaling regulates lung alveolar progenitor cell proliferation and differentiation. Chung *et al*. reported that BMP signaling was active in AT2 in the steady-state, transiently declined post-pneumonectomy in association with the up-regulation of Gremlin1 and other BMP antagonists— and is restored during differentiation of AT2 to AT1 (16).

We show evidence that specific transient overexpression of Gremlin suppresses BMP signaling on day 7 to allow AT2 progenitor activation and subsequent proliferation. Subsequent AT2 proliferation slows down on day 14 evidently through SP-C/Aqp5 ratio being closer to baseline, and BMP signaling is restored closer to baseline. On day 28, we show similar changes but on a higher level of magnitude in comparison to day 14. Mechanistically, it is plausible that the return of BMP signaling and restoration of nearly normal levels of SP-C/Aqp5 between day 7 and day 14 is a result of the large increase in BMP2 production from newly proliferated AT2 cells on day 7. Through scRNA-seq and immunofluorescence, we show that AT2 cells are a major cellular source for BMP2 in the lung. We speculate that this is partly a compensatory effect in which the additional AT2 cells produce BMP2 and 4 ligand at a rate that can overcome the antagonistic effects of Gremlin and restore BMP signaling evident on day 14 and day 28 and drive AT2-AT1 differentiation. The exact amount of BMP ligand needed to overcome Gremlin, however, was not investigated. In the bleomycin-induced lung injury model, the level of Gremlin is indeed upregulated in the early stage (day 7) and normalized in the middle to late stage (day 14 to 28). This mechanistic insight gives us a novel target to look at in the modulation of the AT2 progenitor cell niche in various diseases.

BMP signaling dynamics are important for other progenitor cells. Tadokoro *et al*. reported that inhibitors of the canonical BMP signaling pathway promote the proliferation of basal cells, which can act as progenitor cells in the proximal airway (26).. Similar to AT2s, BMP signaling in the airway normally restricts the proliferation of basal cells at a steady-state, and the transient upregulation of an endogenous BMP antagonist, follistatin, relieves this inhibition during repair Interestingly, from our scRNA-seq data, BMP2 appears to be highly expressed in basal cell along with AT2 cells. BMP signaling and its target genes have also been shown to regulate hair follicle progenitor cell lineages (27). BMP signaling is needed for hair follicle progenitor cell quiescence and to promote differentiation along different lineages as the hair cycle progresses. Taken together, BMP signal dynamics might be one of the common regulators for progenitor cells in multiple organs, not just lungs. Under steady state, BMP signaling is active and prevents uncontrolled progenitor cell proliferation. Under certain situations like injury when progenitor cell proliferation is needed for repairing, BMP signaling is suppressed via increased expression of intrinsic BMP antagonists, which is then restored to halt progenitor cell proliferation and subsequently promote progenitor cell differentiation.

Walsh *et al*. reported a Gremlin promoter contains an NFκB binding site (19). Specifically, two CpG islands contain 6 conserved motifs, 5 of which are NFκB related, and one of them being MA0105, a p50 subunit of NFκB. This suggests a possible regulatory mechanism that pro-inflammatory cytokines may induce Gremlin expression. In the present study, we show evidence that IL-1β induces Gremlin expression in human fibroblasts even at extremely low concentrations in primary culture. Furthermore, given the strong effect of a stiff microenvironment on *GREM1* gene expression in fibroblasts, we speculate that there is likely a strong role of extracellular matrix-mediated mechanotransduction in upregulating Gremlin expression in fibroblasts, and that this may be the difference that sustains Gremlin expression in a disease such as IPF. We propose a positive feedback loop of increasing lung stiffness that activates fibroblasts, continuously upregulating Gremlin and sustaining BMP signaling suppression, potentially leading to AT2 senescence from prolonged proliferation.

It should be noted that there are several intrinsic BMP antagonists. BMP antagonists are classified into three subfamilies based on the size of the cysteine-knot: the DAN family (including Gremlin), twisted gastrulation, and chordin and noggin (28). These BMP antagonists were suggested to have different roles in different organs with varying expression patterns and affinity to BMPs (28). Follistatin, follistatin-like 1, and Gremlin 1 are upregulated in Pdgfr*α*+ mesenchymal cells during alveolar regrowth in pneumonectomy (16), while Gremlin 2 is preferentially expressed in mesenchymal alveolar niche cells (29), suggesting the type of BMP antagonists used in alveolar regeneration may be context-dependent (the type of injury). It would be interesting to investigate the differential expression of BMP antagonists in different types of lung injury.

In summary, we provide novel insight into adult lung wound healing by which Gremlin secreted from macrophages and fibroblasts mediates AT2 proliferation and subsequent differentiation into AT1 through initial suppression of BMP signaling on AT2s to allow AT2-progenitor activity (Summarized in Figure 8). It may have therapeutic implications for several lung injury disorders, including viral/bacterial pneumonia, acute respiratory distress syndrome (ARDS), and emphysema, by enabling interventions that promote proper alveolar epithelial regeneration to re-constitute intact lung architecture.

**Figure 8.**
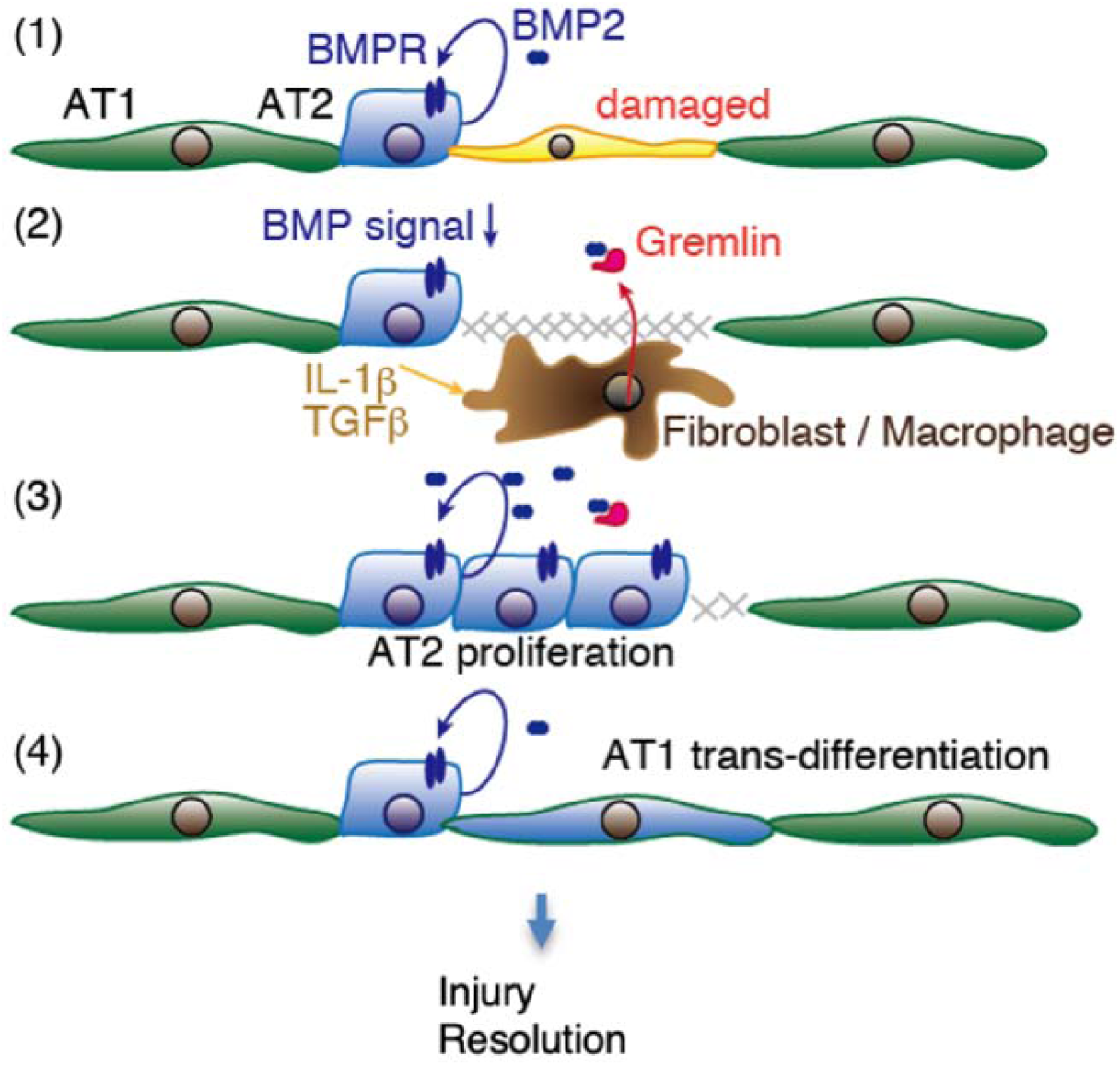
Diagram showing proposed mechanisms by which Gremlin mediates AT2 proliferation and subsequent AT1 differentiation during alveolar injury repair. Once alveolar epithelial cells are damaged, fibroblasts/macrophages migrate around the damaged site. IL-1*β* or TGF*β* stimulates Gremlin production from migrated fibroblasts/macrophages. Produced Gremlin suppresses BMP signaling in AT2s. Suppressed BMP signaling induces AT2 proliferation, and increased AT2s produce BMPs, which overcome Gremlin inhibition. Once BMP signaling is normalized, AT2 proliferation ceases and differentiation from AT2 to AT1 occurs, resulting in wound healing. On the contrary, repeated injury induces fibroblast accumulation and myofibroblast differentiation. These myofibroblasts produce aberrant Gremlin production, which results in sustained BMP signaling suppression in AT2, leading to AT2 dysregulation and fibrosis.

## Acknowledgments

We thank Ms. Fuqin Duan for her technical assistance.

## Notes

### Competing Interest Statement

T. Yanagihara was funded by the Uehara Memorial Foundation Research Fellowship and Mitacs Canada, and the research institute of St Joseph's Hospital, Hamilton, ON, Canada (Post-doctoral Fellowship Award). K. Tsubouchi was funded by the Uehara Memorial Foundation Research Fellowship. K. Ask reports grants and personal fees from Boehringer Ingelheim, grants from Canadian Pulmonary Fibrosis Foundation, Synairgen, Alkermes, GlaxoSmitheKline, Pharmaxis, Unity, Avalyn, Canadian Institutes of Health Research, Ceapro, Pieris, outside the submitted work. M. Kolb reports grants from the Canadian Institute for Health Research and grants/ personal fees from Roche, Boehringer Ingelheim, Prometic, Respivert, Alkermes, and Pharmaxis and personal fees from Genoa.

